# Stationary and germ layer-specific cellular flows shape the zebrafish gastrula

**DOI:** 10.1101/2025.11.10.687746

**Authors:** Susan Wopat, Pieter Derksen, Vishank Jain-Sharma, Gary Han, Nikolas Claussen, Sebastian Streichan

**Affiliations:** Department of Physics, University of California, Santa Barbara, Santa Barbara, California 93106, USA; Department of Cell Biology, Duke University School of Medicine, Durham, North Carolina 27710, USA; Princeton Center for Theoretical Science, Princeton University, Princeton, New Jersey 08544, USA

## Abstract

During gastrulation, a sequence of complex processes transforms the blastula into a multi-layered embryo. Fixed sample analysis has revealed much about how genetic signaling cascades determine the major body axes and organize cell fate patterns into the germ layers: ectoderm, mesoderm, and endoderm. *In toto* live imaging of vertebrate development highlights that embryogenesis is a dynamic process that involves on the order of ten thousand cells, but hurdles related to data handling have hampered quantitative analysis. Therefore, our understanding of the rich physical aspects of multilayered tissue reconfigurations remains incomplete. Here, we reveal that modules of stationary and germ-layer-specific tissue flows shape the zebrafish gastrula. We combine *in toto* live imaging with tissue-specific markers and image analysis to reveal the global shape of the enveloping layer (EVL), epiblast, and mesoderm over time. A user-friendly tissue cartography pipeline based on the Blender 3D software moves into the reference frame of individual tissue layers. We find distinct tissue flow patterns in the enveloping layer (EVL), epiblast, and mesoderm, respectively. The instantaneous tissue flow of these germ layers is organized in a temporal sequence of hours-long, constant flow patterns. This suggests that a sequence of stationary tissue flow modules transports cells to their destination during gastrulation. Mathematical decomposition suggests that epiblast flow is strongly influenced by a superposition of rotational flow in the mesoderm, and divergent flow in the EVL. Molecular and cellular complexity notwithstanding, these results hint at surprisingly, tractable physical processes that underlie vertebrate gastrulation, and set the stage for investigations of how morphogens orchestrate dynamics.

## Introduction

Vertebrate gastrulation organizes the cells of the blastula into germ layers and shapes the blueprint of the body [1, 2, 3]. Traditional and modern approaches to fate mapping in zebrafish reveal distinct biochemical, or transcriptional, signatures of cells that define germ layers [4, 5, 6]. Spatially, lineage tracing has revealed which individual cells contribute to the germ layers [1, 7], and live imaging has also demonstrated migratory motions associated with distinct biochemical markers [8, 9]. Individual cell trajectories are complex [3, 8], pointing to rich physics that shapes the embryo. Rather than analyzing individual cell movements, we take on a germ layer-specific perspective. This view of the embryo has potential to be beneficial when individual germ layers organize into continuous surfaces.

In fish, the initial configuration of germ layers begins when mesendoderm cells differentiate around the margin of the animal cap [10, 11]. This spatial organization defines the starting point of gastrulation and indicates that the problem of shaping the embryo may be broken into two parts: deformation of each germ layer, and cell motion within it. Different layers might execute different shape change programs. Therefore, within each of these dynamic surfaces, we study cell trajectories to understand the programming of tissue shape change from the cellular level. Due to their complexity, it is beneficial to break cell trajectories down into small time units [12, 13]. The resulting *tissue flow field* defines the speed and direction of instantaneous cell motion.

Cells are the active agents that generate forces to drive tissue flows [13]. In many systems, such as quail, chick, and *Drosophila*, anisotropic stresses [12, 14, 15] underlie axis elongation flows. A perspective of fate determinants and lineage tracing suggests that flow is orchestrated by gene expression patterns that move with cells as they rearrange to form the gastrula. In this case, kinematics would be well-described in the co-moving frame of the tissue: a cell’s motion in the flow would only depend on its initial position or that relative to its neighbors, not its instantaneous position on the embryo as a whole. But a priori, it is unclear if transcriptional profiles of cells alone direct their movements without the need for external inputs. In fact, physics-based arguments suggest that systems in which driving cues are advected by tissue flow are often unstable because a small perturbation in the flow can “derail” the guiding cues [16]. Diffusible ligands, such as BMP, provide a spatial pattern that is defined with respect to the embryo as a whole, rather than advected with tissue flows. BMP might play a direct role in organizing motion [17, 18, 19]. In this case, flow would be best described in the embryo frame: a cell’s motion would depend on its absolute position on the embryo. We use tools such as light sheet microscopy and tissue cartography within the 3D software Blender to record and analyze imaging data to settle this question [20, 21, 22]. Our results reveal that tissue flows in the zebrafish embryo are unexpectedly simple in the embryo frame and organized into discrete temporal modules, rather than fluctuating over time. Thus, the direction of cell motion is determined by the spatial position of the cell in terms of longitude and latitude within the associated germ layer. This suggests that the dynamics of morphogenesis unfold in embryo frame.

## Results

### Quantitative imaging pipeline for unconstrained zebrafish embryogenesis

A key difficulty of *in toto* imaging is accommodating large-scale morphogenesis while acquiring a stable, long-term recording of embryonic development. We developed a mounting method for zebrafish embryos that supports tailbud outgrowth during long-term Multiview Selective Plane Illumination Microscopy (MuVi SPIM) imaging. Instead of embedding embryos directly in agarose or methylcellulose, we used a machined mold to form an agarose column with a central conical cavity (**Fig. 1A**). Dechorionated embryos were positioned within the cavity and immersed in egg water within the imaging chamber (**Fig. 1C**). The conical geometry centers embryos automatically, eliminating common off-axis mounting artifacts associated with capillary plungers. This approach enables continuous imaging of the embryo starting at 6 hours post fertilization (hpf) to 24 hpf without detectable developmental defects (**Fig. 1B**).

**Figure 1:**
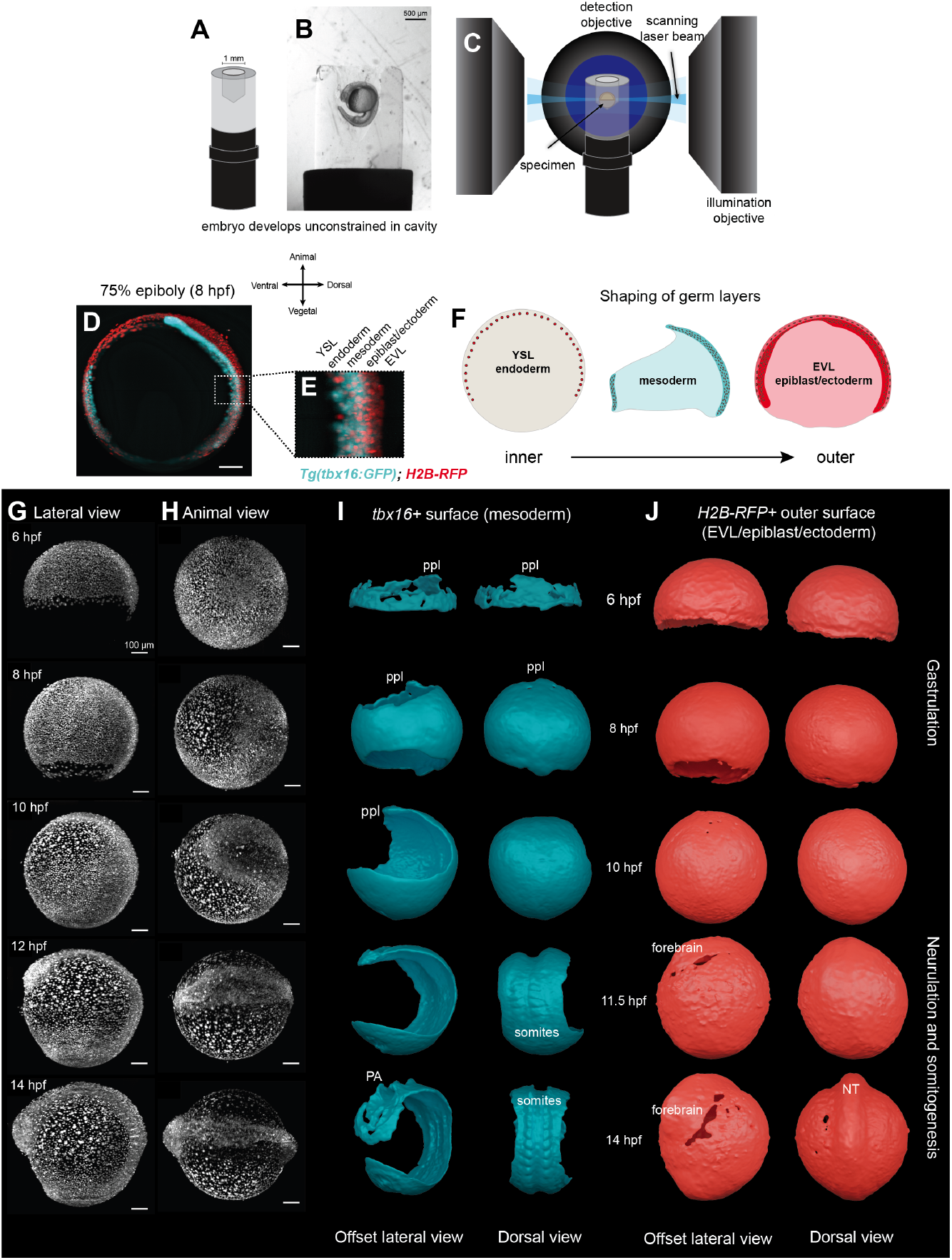
Unconstrained *in toto* imaging of zebrafish development enables extraction of tissue surfaces and visualization of distinct tissue morphologies. **(A)** Schematic of the agarose cavity for the embryo. **(B)** 24 hpf embryo with tail outgrowth following long-term mounting in cavity. **(C)** MuVI SPIM acquisition of embryo developing in central cavity. **(D)** Optical slice of an embryo (8 hpf) imaged with both the nuclear marker, *H2B-RFP* and the mesodermal marker, *Tg(tbx16-eGFP)* shows morphology of layered tissues. Scale bar is 100 *µ*m. **(E)** Inset highlights the discrete mesodermal layer (cyan), while all tissues are labeled with *H2B-RFP*. **(F)** Schematic representing germ layer surfaces and shapes at 8 hpf. **(G)** Lateral view of an embryo developing in cavity (6, 8, 10, 12, and 14 hpf). *H2B-RFP* labels all nuclei. **(H)** Animal view of same embryo. **(I)** Offset lateral and dorsal views displaying extracted isosurfaces of the tbx16+ tissue layer at 6, 8, 10, 11.5, and 14 hpf. Mesoderm-derived tissues such as the prechordal plate (ppl), somites, and pharyngeal arches (PA) are visible. **(J)** Offset lateral and dorsal views displaying extracted EVL/epiblast/ectoderm isosurfaces at 6, 8, 10, 11.5, and 14 hpf. The neural tube (NT) and forebrain emerge at 11.5 hpf and become increasingly apparent at 14hpf. NT is clearly visible at 14 hpf. Lower nuclear densities at later stages account for holes in the surface mesh where the epidermis covers the yolk.

### *In toto* imaging of zebrafish gastrulation with germ-layer markers

Using this approach, we acquired time-lapse recordings of zebrafish embryos injected with *H2B-RFP* mRNA to label all nuclei along with a mesodermal reporter *Tg(tbx16:eGFP)* [23] (**Fig. 1G and 1H**). Eight views of both channels were collected every two minutes from 6–14 hpf and fused using the Multiview Reconstruction plugin in ImageJ [24] to generate 3D volumes with isotropic resolution. Although our mounting approach permits imaging to 24 hpf without issue, we focused on the time windows of gastrulation and early neurulation for this study.

With a complete view of the entire embryo, we first asked how distinct extraembryonic and embryonic tissues shape themselves to collectively build the gastrula. Via image segmentation, we extracted *tbx16*-positive cells to distinguish mesoderm progenitor cells from the other germ layers and extraembryonic tissues, including the yolk syncytial layer (YSL) and enveloping layer (EVL) (**Fig. 1D, 1E and 1F**). Three-dimensional renderings revealed dramatic mesoderm reshaping over gastrulation, including the dorsal accumulation of tissue to form the midline. At later stages, somites and pharyngeal arches also become apparent (**Fig. 1I**). It is important to note that the *tbx16:eGFP* transgenic reporter does not label chordamesoderm, meaning the surfaces we generated primarily represent the prechordal plate and trunk mesoderm [25]. We then specifically segment the *H2B-RFP* nuclear signal in cells above the *tbx16*-positive signal to obtain the epiblast, or presumptive ectoderm surface, along with the thin outer EVL extraembryonic tissue. Unlike the mesoderm, dramatic shape changes in the epiblast/ectoderm (EpE) primarily arise after gastrulation, when formation of the forebrain and neural tube begin to build (**Fig. 1J**). Tissue-specific segmentation reveals that each germ layer adopts a distinct geometry during gastrulation. By decomposing the embryo’s global remodeling into distinct tissue layers, and quantifying their shape and dynamics as digital surfaces, this approach provides a clear view of how individual tissue morphologies emerge.

### Mapping 3D germ-layer surfaces in 2D

To measure the collective cellular dynamics underlying these shape changes, we used image processing approaches to examine cell movements within different tissue layers. Conventional 3D nuclear tracking of *in toto* embryogenesis becomes unreliable once the embryo comprises thousands of cells. Furthermore, 3D tracks offer information on individual cells, but fail to provide insights into the collective, coherent motions of large groups of cells, which is essential for understanding tissue-level deformation. Thus, to access tissue movements, we converted our 3D volumetric data into 2D surface projections of individual tissue layers (**Fig. 2**), akin to 2D cartographic projections of the globe. This successfully produced simplified 2D surface maps of mesodermal and epiblast/ectodermal/EVL nuclei from the more complex 3D shapes segmented from the *tbx16-eGFP* and *H2B-RFP* signals, respectively (**Fig. 2H-2H”**). Importantly, this cartographic framework preserves tissue coordinates, enabling re-mapping of projections onto the three-dimensional embryo (**Fig. 2G-2G**“) [21, 22]. Furthermore, different timepoints are registered into a common cartographic reference frame, allowing analysis of tissue dynamics and cell flow.

**Figure 2:**
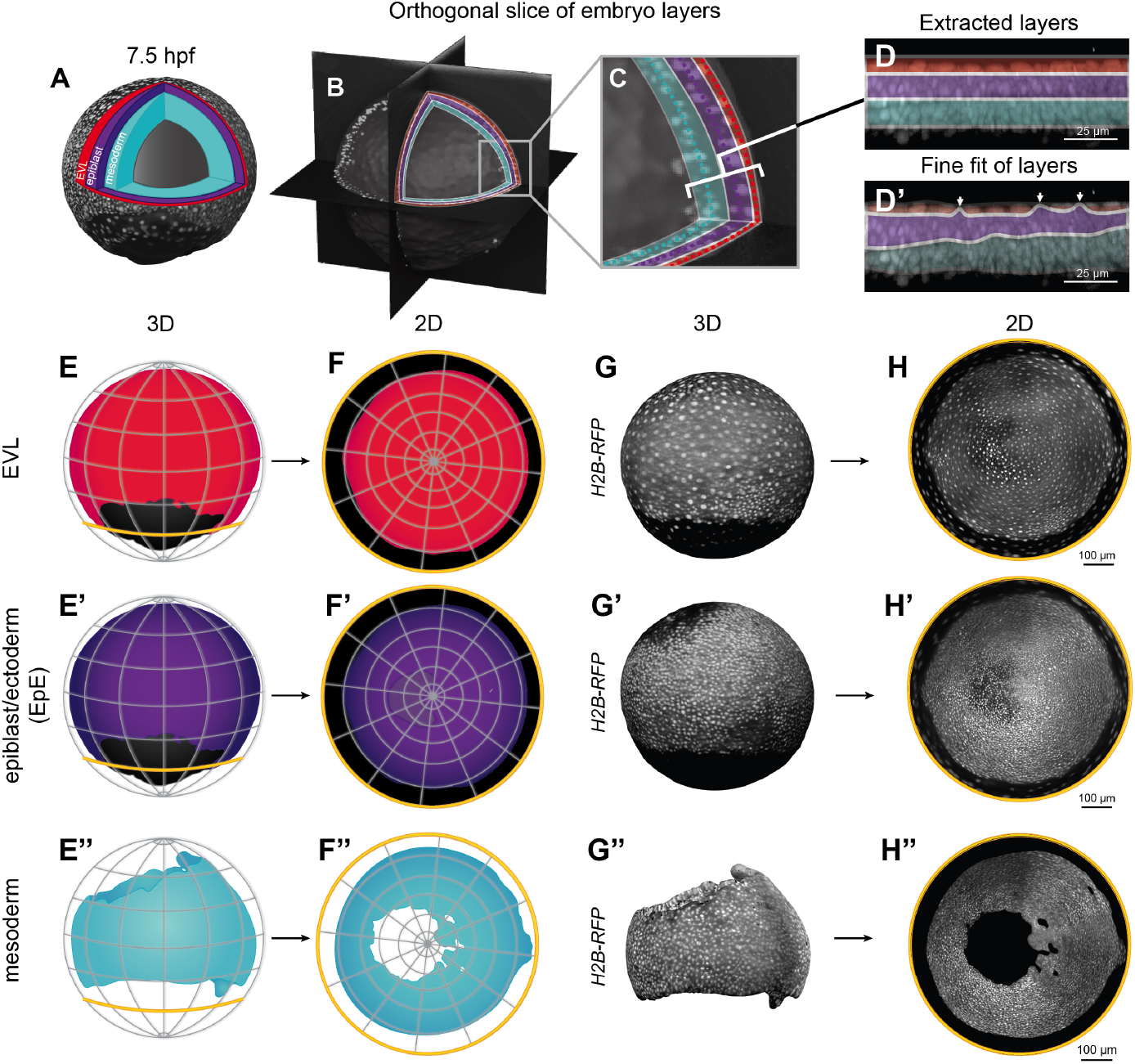
Visualizing and mapping the embryo layer by layer. **(A)** A 3D surface mesh of a 7.5 hpf embryo of *H2B-RFP* expression. Overlaying schematic details how tissue layers can be “peeled off” and extracted separately. We extract the outermost tissue EVL (red), epiblast (purple), and mesoderm (cyan) layers. **(B)** Color-shading of orthogonal sections showing actual tissue layering of the EVL (red), EpE (purple), and mesoderm (cyan). **(C)** Inset highlighting varying thickness within layers **(D-D’)** Fine fit correction on extracted layers ensures precise segregation of tissues for 2D mapping. White arrows point to areas of the EpE layer that must be removed to finely extract the thin EVL **(E-E” ‘)**. Schematics of the 3D surfaces for the EVL, EpE, and mesoderm. **(F-F” ‘)** Schematic of the corresponding 2D polar maps generated from the 3D surfaces of each tissue layer. **(G-G”‘)** *H2B-RFP* nuclei expression in each tissue layer (EVL, EpE, mesoderm) represented on 3D surface meshes. **(H-H” ‘)** Corresponding 2D polar maps of *H2B-RFP* nuclei extracted from the 3D surfaces of each tissue layer (EVL, EpE, and mesoderm). 2D maps represent max intensity projections of all layers.

More precisely, 2D projections of tissue-specific maps for the mesoderm, EpE, and the outermost EVL tissue are first coarsely generated by “peeling off” the layers of the embryo surface separately (**Fig. 2A-F** from the outside in, i.e. along the local surface normal) [21]. While this study focuses on the three outermost tissue layers, the approach readily extends to the innermost tissues — the endoderm and YSL cells — with the use of an endoderm transgenic marker. To finely fit our layers to the actual tissue shapes, such as the EVL, a very thin tissue that overlays the EpE, we performed a “fine fit” that adjusts for small changes in the surface, to prevent off-target extraction of the EpE nuclei (**Fig. 2C-D’**). In total, 60 timepoints (6–8 hpf) were processed to produce 2D layer-specific projections for the mesoderm, EpE, and EVL (**Fig. 2H-H”**, **Methods**). Importantly, surface extraction can be applied to any transgenic line — not just nuclear lines — to visualize global levels of gene expression, localization of cytoskeletal machinery, and cell morphology dynamics. The Blender tissue cartography pipeline (**Methods**, [22]) enhances surface extraction, visualization, and mapping to account for fine surface features. By enabling the separation of very thin layers of the zebrafish gastrula for the first time, these tissues become accessible for *in toto* analysis. We then generated a max intensity projection to create a single, coherent map from 3D to 2D for timepoints 6-8 hpf, so that tissue dynamics could be visualized and analyzed in 2D. We registered the different timepoints into a common reference frame. Note that this will also be useful for comparing multiple movies and experimental conditions.

### Germ-layer-specific flow fields during gastrulation

To quantify tissue movements between 6–8 hpf, we used particle image velocimetry (PIV) instead of tracking individual nuclei. PIV is widely used to analyze tissue flows [26], and can accurately extract motion patterns in different developmental contexts [13]. Here, we use PIV to analyze nuclear displacement across the developing mesoderm, EpE, and EVL tissue layers. By integrating the PIV flow fields, “virtual” cell trajectories can be reconstructed (**Fig. 3A-C**).

**Figure 3:**
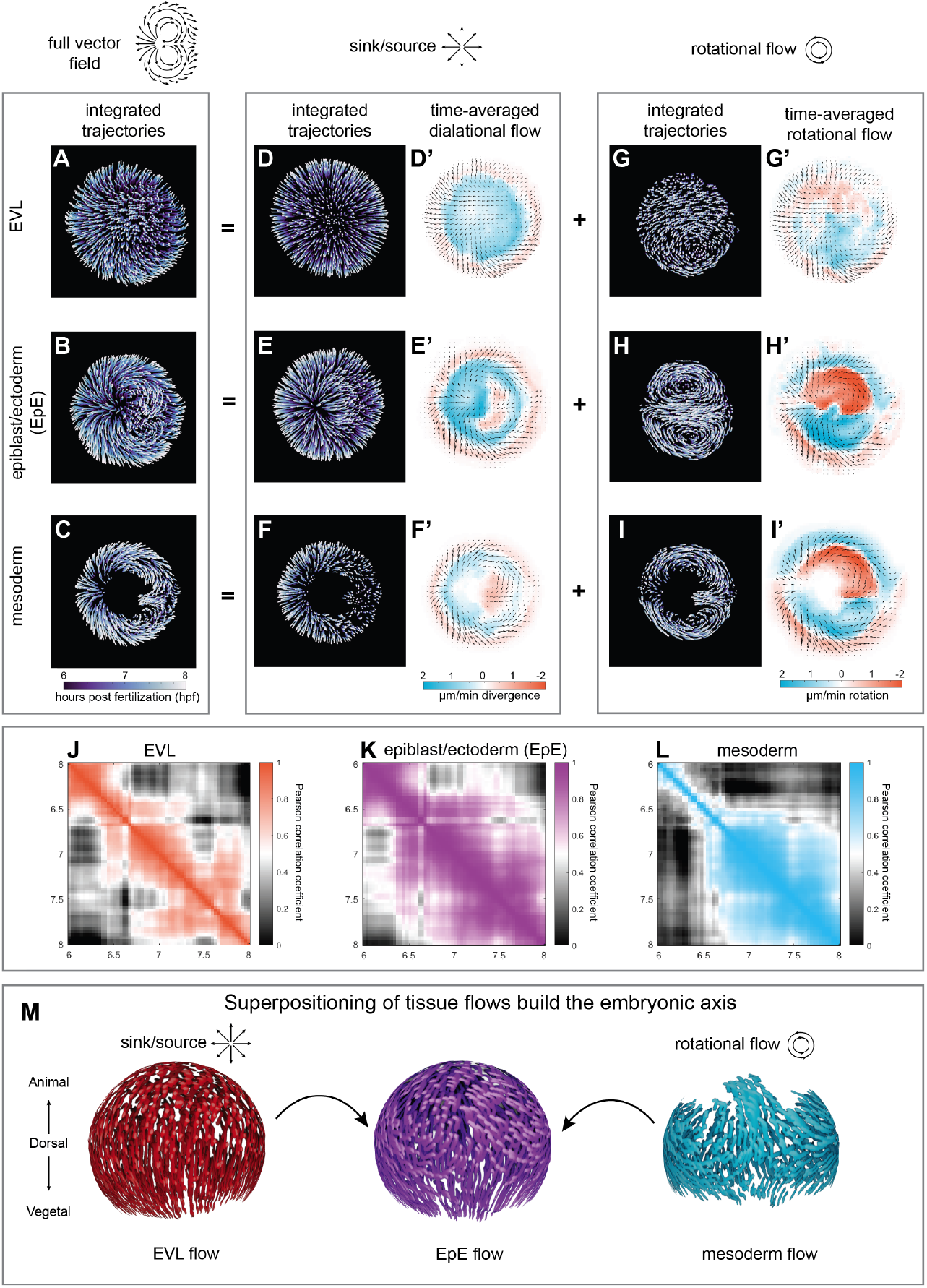
Dynamics of shape change in EVL, ectoderm, and mesoderm. **(A-C)** Integrated trajectories of tissue motion from 6-8 hpf in (A) EVL, (B) EpE, and (C) mesoderm. **(D-F)** Integrated trajectories of the source-sink flow component in EVL, (E) EpE, and (F) mesoderm. **(D’-F’)** Magnitude of the dilational component with positive and negative signs corresponding to sources or sinks in the divergence, in (D’) EVL, (E’) EpE, and (F’) mesoderm. Average of full vector field is overlaid. **(G-I)** Integrated trajectories from 6-8 hpf of rotational component in (G) EVL, (H) EpE, and (I) mesoderm. **(G’-I’)** Magnitude of the rotational component with positive and negative signs corresponding to clockwise or anti-clockwise movement circulation in (G’) EVL, (H’) EpE, and (I’) mesoderm. Average of full vector field is overlaid. **(J-L)** Autocorrelation matrix of flow fields from 6-8 hpf in the (J) EVL, (K) EpE, and (L) mesoderm. **(M)** Renderings of 3D flow fields show dominant sink/source flow in the EVL (red) and rotational flow mesoderm (cyan). These flow features are represented in the EpE layer (purple), suggesting that flows in the EVL and mesoderm influence the kinematics of the ectoderm.

In the EVL, cells primarily expanded toward the vegetal pole. However, we also observed that on the dorsal side, nuclei moved slightly toward the animal pole at later gastrulation time points (**Fig.3A**). Both EpE and mesodermal layers exhibited bilaterally symmetric vortices converging toward the presumptive midline, characteristic of convergent extension movements previously described in the avian embryo [12, 15] (**Fig. 3B and 3C**). In the mesoderm, two large vortices with lateral fixed points (positions where cells do not move) likely arise from the combination of organizer, or embryonic shield, ingression and the dorsal-directed migration of ventrolateral cells. In the EpE, vortical flow patterns may originate from intrinsic dynamics within the tissue or via external forces transmitted through mesoderm. However, previous findings support the latter interpretation. In *one-eyed pinhead* mutants that lack mesoderm formation, vortical motion in the overlying EpE is markedly reduced, indicating that EpE cell movements depend on frictional coupling between the two layers [9].

### Decomposing flow fields into separate components reveals simple physical principles of axis formation

These observed qualitative differences between flow fields led us to ask how these flow motifs within individual layers collectively shape the embryonic midline. To uncover the underlying organizational principles, we decomposed the velocity field into dilational and rotational components (**Fig. 3D-I**). Mathematically, the two components correspond to a curl-free and a divergence-free vector field, respectively (**Fig. 3D-I**). In the EVL, integrated trajectories and a spatial analysis of the source/sink component show that tissue movement is strongest at the margin (**Fig. 3D-D’**), corresponding to the expansion during epiboly. The EpE also has a very strong source/sink flow, with a fixed point evident in the trajectories that appears to correspond with the emerging midline (**Fig. 3E-E’**). In the mesoderm, tissue expansion is primarily localized to the ventral side, but is weaker on the dorsal side (**Fig. 3F-F’**). On the other hand, the rotational component is minimal in the EVL, while it is much higher in the EpE and mesoderm tissues (**Fig. 3G-I’**). The rotational component in the EpE clearly shows two dominant vortices that define the presumptive midline (**Fig. 3H-H’**). This flow pattern is also reflected in the mesoderm (**Fig. 3I-I’**).

We next generated autocorrelations of the flow fields within each layer, asking whether flows are approximately stationary over time (similar approach to [27]). We notice a relatively high degree of correlation within the EVL flow over an evolving 30 min time window (**Fig. 3J**). However, in the EpE, we see two modules of high similarity, indicating that the flow abruptly changes character (**Fig. 3K**). Interestingly, the timing of this switch aligns with the emergence of a flow module in the mesoderm, which begins at approximately 6.75 hpf (**Fig. 3L**). Based on our component analysis and spatial profiling of the mesoderm, this block corresponds with the emergence of the two dominant vortices (**Fig. 3I**). Thus, this result suggests that the rotational flow of the mesoderm triggers a rotational flow in the EpE layer.

Together, these results reveal that axis formation emerges from a simple set of conserved kinematic motifs— dual vortical rotation and directional expansion—coordinated across germ layers (**Fig. 3M**). By resolving seemingly complex flow fields into a small number of components, we provide a physical framework for examining dynamic tissue behaviors in a variety of contexts. More specifically, this generates a unified morphogenesis framework that can generate quantitative comparisons across multiple perturbations without the need for complicated analyses or cell tracking. This method sets the stage for linking morphogenetic signaling to the underlying forces that control large-scale embryonic symmetry and body-plan organization.

## Discussion

Vertebrate embryogenesis involves elaborate biochemical signaling pathways that act in tens of thousands of cells to establish the blueprint for intricate shapes [3, 6, 8, 28]. Here, we find germ-layer-specific flows are stationary for extended periods of time and shared flow motifs suggest mechanical interaction across layers, as has been described before [9, 29]. This quantitative analysis of kinematics during gastrulation from a germ-layer perspective suggests an astoundingly simple view. In contrast, minimal *in vitro* models for active, molecular-motor-driven flows, which comprise a small number of reconstituted cytoskeletal components, are typically complex and chaotic [30, 31]. Therefore, the relative simplicity of flow patterns in vertebrate gastrulation suggests that development evolved molecular complexity to robust physics [14, 15, 32]. How can stationary patterns of flow emerge when cells carry the active force-generating machinery with them as they move? For invertebrates, tissue dynamics have been suggested to feedback to the organization of the cytoskeleton [33, 34, 35]. It will be exciting to test if similar mechanical mechanisms exist in zebrafish embryos. Additionally, zebrafish’s genetic tractability also presents an ideal system to further explore the interplay of Mangold organizer-signaling gradients and feedback.

## Materials and Methods

### Fish stocks

Adult zebrafish of the TU and AB strains were housed and bred using standard conditions. All experiments were performed in accordance with institutional approval from the University of California, Santa Barbara. Previously published transgenic lines used for this study include *Tg(tbx16:GFP)*uaa3 [23].

### Light sheet microscopy

For generating *in toto* movies of zebrafish development, we use a custom-built Multiview Selective Plane Illumination Microscope [36] in a scatter-reducing, or confocal, imaging mode [20]. Briefly, the microscope design involves duplicate pairs of orthogonally arranged illumination and detection arms, such that illumination and detection objectives face one another. For illumination, we utilize a custom-built laser combiner that houses continuous wave laser lines, including 488 and 561 nm, all OBIS LX, Coherent Inc. These laser lines always operate at the same set output power for each acquisition. Stability of the laser beams was assessed with an optical power meter from Thorlabs (PMD 100D with S121C). These laser lines were combined using dichroic mirrors on kinematic mirror mounts. To duplicate the light path for feeding the two illumination arms, a broadband beam splitter from Omega Optical Inc., evenly splits beams for 488 and 561 nm. Both light paths are identical and consist of a pair of kinematic mirrors for alignment purposes, a galvanometric mirror (Cambridge technology), a scan lens (Sill Optics), a Tube lens (200 mm focal length), and a water-dipping objective (CFI Plan Fluor 10x, NA 0.3, Nikon). The detection involves a water-dipping objective (CFI75 LWD 16x, NA 0.8 Nikon), a filter wheel (Lambda, Sutter Instruments) with emission filters (FF01-542/27-25, FF01-609/62-25, BLP01-568R-25, BLP01-664R-25, all Semrock), a tube lens (200 mm focal length), and an sCMOS camera (Hamamatsu ORCA-Flash 4.0 V3).

For optimal image quality, we reduced optical scattering, following the strategy outlined in Ref. [20]. Briefly, we employ a National Instruments Multi-function I/O card (PCI-6229) to generate a sequence of electronic signals to synchronize the galvanometric mirror phase with the start of the image acquisition and stage movements. The camera operates in a light-sheet mode readout, which is set to the maximum speed for the acquisition front. The width of exposing pixels around this form was set to 52 pixels (corresponding to an effective 13.4 µm) in all experiments.

Optical sectioning involved electronically controlled stages, all from Physik Instrumente GmbH and Co.Kg. A translation stage (linear piezo stage P-629.1 cd with E-753 controller), a rotational piezo stage (U-628.03 with C-867 controller), and a linear actuator (M-231.17 with C-863 controller). The electronic acquisition was controlled through Micro-Manager [37].

### Image acquisition

Embryos are manually dechorionated using forceps and mounted in conical-shaped cavities created in agarose pillars. The embryo is then imaged simultaneously from two detection objectives with z-step of 2.0 mm to produce two volumetric stacks of the entire embryo from opposite sides. Then, the embryo serially rotates 45 degrees to capture three additional positions, thereby producing eight views per time point. Total imaging duration per time point is ∼50 s, allowing for temporal resolution of 120-300 s.

### Image fusion

Fluorescent beads (Fluoresbrite multifluorescent 0.5-*µ*m beads 24054, Polysciences Inc.) are diluted 1:1000 in 2% low-melting point agarose in egg water solution to make agarose pillars. The fluorescent beads serve as fiducial markers for registering views with interest point detection using the Fiji plugin, “Multiview Reconstruction” [24]. “Difference of Gaussian” interest point detection identifies bead locations in all views for all time points. Interest point positions are then correlated using the fast 3D geometric hashing (rotation invariant) algorithm with reasonable global all-to-all time point matching (sliding window of five time-points). An affine transformation regularized to a rigid model using a lambda value of 0.10 is used to register views. Lastly, 3D stacks of all the views are fused or deconvolved with an efficient Bayesian iteration. The PSF estimation is extracted from the beads to deconvolve the resulting image, which has an isotropic resolution of 0.4092*µ*m.

### Pullback generation: 3D analysis and cartographic projection

3D data was analyzed using the `blender - tissue - cartography` (`btc`) workflow described in Ref. [22]. In brief, this workflow comprises three steps. First, the outline of embryos was segmented out of volumetric data using `ilastik` [38]. Second, volumetric data and segmentations were loaded into the 3D software `Blender` using the `btc` add-on, converting segmentations into polygonal meshes of the embryo surface. Meshes and 3D data were visualized in `Blender`, using `btc` to project the 3D data onto the mesh surface. Third, meshes were cartographically unwrapped to the plane using `Blender`’s UV mapping editor, and `btc` was used to project image intensities in selected layers parallel to the embryo surface to 2D.

To analyze time-lapse data, the surface meshes for all time points were loaded into a single `Blender` file. A “reference” sphere, equipped with a desired cartographic map (e.g a polar projection), was aligned around the meshes (rotated so the poles of the reference sphere matched the animal and vegetal poles). To project a mesh point to 2D, it was radially traced to the reference sphere and then mapped to 2D using the sphere’s cartographic map (see **Fig. 2E-E”**). In this way, all time points are projected to 2D within a common reference frame, allowing, for instance, analysis of cell motion flow via PIV. In all quantitative analyses, `btc` was used to mathematically correct for cartographic distortion.

### Particle image velocimetry (PIV)

For generating tissue velocities on the surface of the embryo we used the Particle Image Velocimetry (PIV) method to compute the instantaneous tangent velocity field of the tissue (similar to [27]). The PIV algorithm divides an image into smaller windows and measures the displacement of each window between subsequent timepoints. Therefore, PIV yields a measure of tissue velocity on a coarse grid, whose spacing equals the window size. After generating 2D cartographic projections of surface tissue movies, we computed PIV in the 2D map space on the vertices of a square lattice with edge length of approximately 2% of the map range in both directions (hence, spanning multiple cell widths). PIV velocity fields were computed for all movie timepoints using a custom MATLAB script. This procedure resulted in a spatially discrete time-varying velocity field 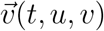 across the lattice grid, with *t* denoting time and (*u, v*) denoting a rectangular parametrization of 2D map space. These coarse-grained velocity fields were then used to perform subsequent flow analysis, including component decomposition and trajectory integration.

Spatial distributions of the magnitude of each of the dilational and rotational components for the different tissue layers were computed from the magnitude of the corresponding velocity fields, with distortions of the velocities due to cartographic projection corrected by re-projecting the vector fields back into 3D. To give these magnitudes a meaningful sign in the context of tissue flows, we calculated the divergence and curl of the total velocity field, and then averaged them over the timerange of interest. Here, positive and negative signs correspond to sources or sinks in the divergence, and clockwise or anti-clockwise movement in the curl, respectively.

### Integrated flow trajectories

To generate flow trajectories for both total and decomposed flow fields, we used a Runge-Kutta 4th order scheme to integrate PIV velocity fields over time. We performed numerical integration on our discrete velocity data by linearly interpolating the velocity field in space and time. To smoothen the trajectories in time, we used a numerical integration timestep with resolution of 0.67 minutes in the Runge-Kutta integration (the value was chosen such that multiple smoothing time steps were integrated within the time resolution of our movies, but its value is arbitrary). For the trajectories shown in **Fig. 3**, all integrations were performed within time interval *T* of 6 hpf to 8 hpf, with the trajectories illustrating the cumulative tissue motion within *T*. To visualize the trajectories, a dense set of points was randomly generated within the 2D space (excluding the regions outside the tissue), and these points were used as initial conditions in all integrations. Resultant trajectory patterns reflected the underlying flow fields, and were insensitive to choice of initial points. Trajectories were parametrized by the fraction *f* of time *t* passed within the overall time interval, given by *f* = *t/T*, and display colors correspond to the value of *f*.

## Acknowledgments

We would like to thank the UCSB Animal Resource Center for fish care. We also thank Stephanie Woo for helpful discussions on the manuscript and experiments as well as members of the Streichan group for thoughtful feedback.

## Contributions

S.W., V.J.-S., P.D. and S.J.S. conceived and designed this project. S.W. performed experiments. S.W., V.J.-S., P.D., G.H., N.C. and S.J.S. performed data post-processing. V.J.-S., S.W., P.D., G.H., N.C. and S.J.S. analyzed the data. P.D., V.J.-S., N.C. and G.H. developed the original image analysis tools. S.W. and S.S. wrote the manuscript with input from all authors.

